# Hepatocyte Circadian Clocks Control Cholesterol Metabolism and Protect From Metabolic Dysfunction-Associated Steatohepatitis (MASH)

**DOI:** 10.1101/2025.08.07.669039

**Authors:** Leonardo Vinicius Monteiro de Assis, Lina Jegodzinski, Julica Inderhees, Sylvia Wowro, Juliana Marques Affonso, Isabel Heyde, Emelie Luise Fischer, Witigo von Schönfels, Andrea Schenk, Florian Roßner, Michael Schupp, Jens U. Marquardt, Münevver Demir, Henrik Oster

**Author notes:** Corresponding authors: Leonardo VM de Assis Henrik Oster.

## Abstract

The circadian clock synchronizes physiological processes with the 24-hour light-dark cycle. Clock disruption contributes to metabolic disorders, including metabolic dysfunction-associated steatohepatitis (MASH). Here, we investigated the role of the hepatocyte clock in MASH using hepatocyte-specific *Bmal1* deletion (Hep-Bmal1KO) mice. Hep-Bmal1KO mice showed faster MASH progression with increased hepatic cholesterol, inflammation, and fibrosis. Transcriptomic and lipidomic analyses revealed dysregulated cholesterol metabolism in Hep-Bmal1KO mice, marked by reduced expression and disrupted rhythmicity of key cholesterol-related genes. Bioinformatic analyses identified *Chrebp* as a potential co-regulator of these transcriptional changes. In an in vitro model with palmitate exposure and gene silencing, we found that *Bmal1*, but not *Chrebp*, regulated cholesterol accumulation, indicating *Bmal1’s* specific role in hepatic cholesterol metabolism. Translating our findings to a human patient cohort revealed a significantly shifted circadian phase, despite no marked effect on hepatic cholesterol levels in the livers of patients with more advanced liver disease (i.e., MASH) compared to simple steatosis. Taken altogether, our findings offer a roadmap to understand the hepatocyte clock’s role in MASH and its potential as a therapeutic target.

## INTRODUCTION

The circadian clock is a highly conserved temporal system that synchronizes biological processes with the Earth’s 24-hour rotation, enabling organisms to anticipate environmental changes and coordinate physiology according to the cyclic demands of their surrounding ^1^. At the molecular level, the circadian system is based on interlocked transcriptional-translational feedback loops (TTFLs) of core clock genes that oscillate throughout the day, regulating the expression of numerous gene programs ^2^. A central clock in the suprachiasmatic nucleus (SCN) coordinates peripheral clocks through multiple pathways, including behavioral (e.g., food intake and activity/rest rhythm), humoral (e.g., cortisol and melatonin), and neuronal signals (e.g., autonomic activity) to align with the external day-night cycle ^3,4^.

The liver functions as a central metabolic hub and is the main site of carbohydrate, lipid, amino acid, bile acid, and xenobiotic metabolism, which are all under the control of the circadian clock ^5–7^. Shift work and other factors, such as daily stress and high-calorie diets, are known to disrupt the circadian clock network, and chronodisruption is a risk factor for the development of Metabolic-Dysfunction-Associated Steatotic Liver Disease (MASLD) ^8,9^. MASLD has become the most common chronic liver disease globally, affecting about 30 % of adults in Western countries ^10^. MASLD is caused by the liver’s inability to metabolize carbohydrates and fatty acids effectively. Additional imbalances, such as in de novo lipogenesis, beta-oxidation, triglyceride secretion via very low-density lipoproteins, endoplasmic reticulum stress, mitochondrial dysfunction, inflammation, insulin resistance, microbiome, and increased fibrogenesis, promote the transition of MASLD to more severe conditions, such as metabolic dysfunction-associated steatohepatitis (MASH), liver fibrosis, or hepatocellular carcinoma (HCC) ^11,12^.

Circadian disruption impacts the liver and contributes to MASLD development. For example, chronic jetlag exposure of wild-type mice leads to steatosis in all animals and HCC in up to 9 % of them. HCC rates are even higher in circadian clock mutant mice because of elevated androstane receptor (*Car*, also known as *Nr1l3*) signaling ^13^. In line with this, mice kept for two weeks on a choline-deficient high-fat diet (cdHFD) develop MASH and display a notable 4-hour phase advance in core clock genes in the liver compared to chow-fed mice, though clock gene amplitudes remain largely unchanged. At the same time, many clock target gene rhythms exhibit substantial changes in phase and amplitude ^14^. Along with liver disease progression – from MASLD to fibrosis – circadian rhythms deteriorate further ^6^.

In this study, we examined the role of the hepatocyte clock in MASH. Targeted *Bmal1* deletion in hepatocytes resulted in significant liver transcriptome rewiring under control conditions, which was potentiated under MASH conditions. The absence of a hepatocyte clock worsened MASH progression, leading to higher levels of hepatic cholesterol, inflammation, and fibrosis. Predictive analyses identified carbohydrate-responsive element-binding protein (CHREBP) as a potential co-regulator of *Bmal1*-driven effects. Notably, genes affected by *Bmal1* knockdown, but not *Chrebp*, were associated with cholesterol but not triglyceride metabolism. Similarly, *Bmal1*-silenced hepatocytes treated with palmitate, an in vitro model of MASLD, showed increased cholesterol levels compared to control cells. In a patient cohort, MASH patients presented with a shifted circadian phase in comparison to patients with simple steatosis.

Taken together, we show a role of the hepatocyte circadian clock function in MASH progression, emphasizing its protective effects as a potential therapeutic target.

## RESULTS

### Hepatocyte-specific *Bmal1* deletion reduces amplitude, advances the phase of circadian gene expression, and disrupts core clock gene rhythms in mouse liver

The Hep-Bmal1KO mouse model, created more than ten years ago ^15^, allows to specifically investigate the influence of hepatocyte clocks on circadian physiology. We collected liver samples from young adult Hep-Bmal1KO mice and age-matched controls (Hep-Bmal1WT) of both sexes every 4 hours over a 24-hour cycle (Figure 1 A). To identify the presence of rhythmicity in mRNA profiles, we employed a suite of analytical tools, including JTK_cycle, Metacycle, CircaN, and DryR, together with a stringent statistical threshold (p < 0.01). 2,756 and 1,327 genes were exclusively rhythmic in livers of Hep-Bmal1WT or Hep-Bmal1KO, respectively, whereas about 50% of the rhythmic transcriptome (3,224 genes) were rhythmic in both groups (robustly rhythmic genes – RRGs) (Figure 1 B, C). Rhythmic genes, identified in at least one group, were then integrated into CircaCompare to identify rhythm parameter changes between genotypes. RRGs showed a general reduction in amplitude and phase advances in Hep-Bmal1KO livers (Figure 1 D). Similarly, a decrease in amplitude and a tendency for a phase advance (p = 0.14) were identified for the core clock genes (Figure 1D). Reduced amplitudes were identified in genes from the positive (*Bmal1(Arntl)*, *Clock*, and *Npas2*), negative (*Cry*s and *Per*s, except for *Per1*), and auxiliary loops (*Nr1d1/2* and *Dbp*) of the clock TTFL (Figure 1 E).

**Figure 1:**
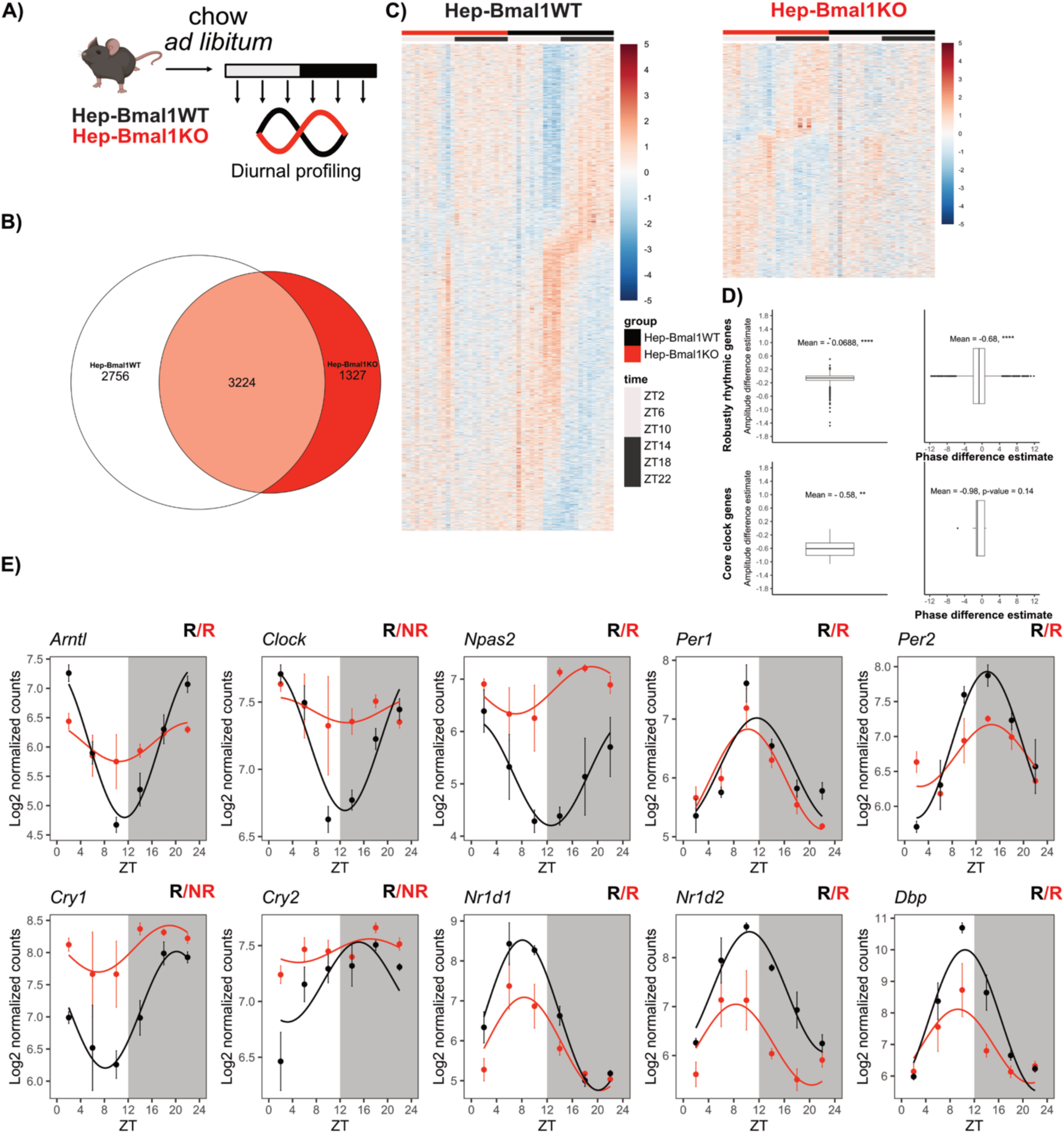
Deletion of *Bmal1* in hepatocytes dampens and shifts the phase of the rhythmic liver transcriptome. A) The design of the in vivo experiment is depicted. B) A Venn diagram shows the number of rhythmic genes in Hep-Bmal1WT and Hep-Bmal1KO mice. C) Heatmap represents the exclusive rhythmic genes of Hep-Bmal1WT (left) and Hep-Bmal1KO mice (right). D) Boxplot shows the difference in amplitude and phase between genotypes for the robustly rhythmic and core clock genes. E) The diurnal profile of core clock genes is depicted. N = 4 per ZT. ‘R’ symbol indicates rhythmicity, while ‘NR’ signifies arrhythmicity detected by CircaCompare.

### Loss of *Bmal1* in hepatocytes dampens mRNA rhythms and lowers overall expression levels of metabolic and transcription factors (TFs)

Detailed rhythm parameter analysis using CircaCompare identified several genes with altered mesor, amplitude, and/or phase (differentially rhythmic genes – DRGs). A total of 2,168, 629, and 450 genes showed changes in mesor, amplitude, or phase, respectively (Figure 2 A; Table S1). In Hep-Bmal1KO livers, genes with increased mesor showed enrichment for biological processes associated with biosynthetic activities, including amino acids, isoprenoids, sterols, and organophosphates. Additionally, enrichment was seen for processes of structural assembly (e.g., ribosome and subunit biogenesis) and DNA repair (e.g., double-strand break repair). DRGs exhibiting reduced mesor were enriched for lipid biosynthesis as well as glycerolipid and carbohydrate metabolism (e.g., monosaccharide metabolic processes). Additional processes associated with energy-associated pathways (e.g., generation of precursor metabolites), membrane transport (e.g., monocarboxylic acid transport), and response to cholesterol were also identified. Genes with a reduction in amplitude were enriched for processes related to circadian rhythms and various metabolism-related processes (e.g., alpha-amino acid, polysaccharide, carboxylic acid, and vitamin B6 metabolism). Conversely, genes that gained amplitude were enriched for immune functions (e.g., including myeloid cell differentiation and the regulation of the adaptive immune response) and some metabolic processes, such as fatty acid and organic acid biosynthesis, in addition to steroid and alcohol metabolism. Among those genes with a phase advance, biological processes related to energy metabolism (e.g., carbohydrate and purine metabolism, and ATP biosynthetic processes) and lipid storage were overrepresented. Conversely, genes with a phase delay were enriched for metabolic processes related to ketones, lipids, fatty acids, and cholesterol metabolism (Figure 2 B; Table S1).

**Figure 2:**
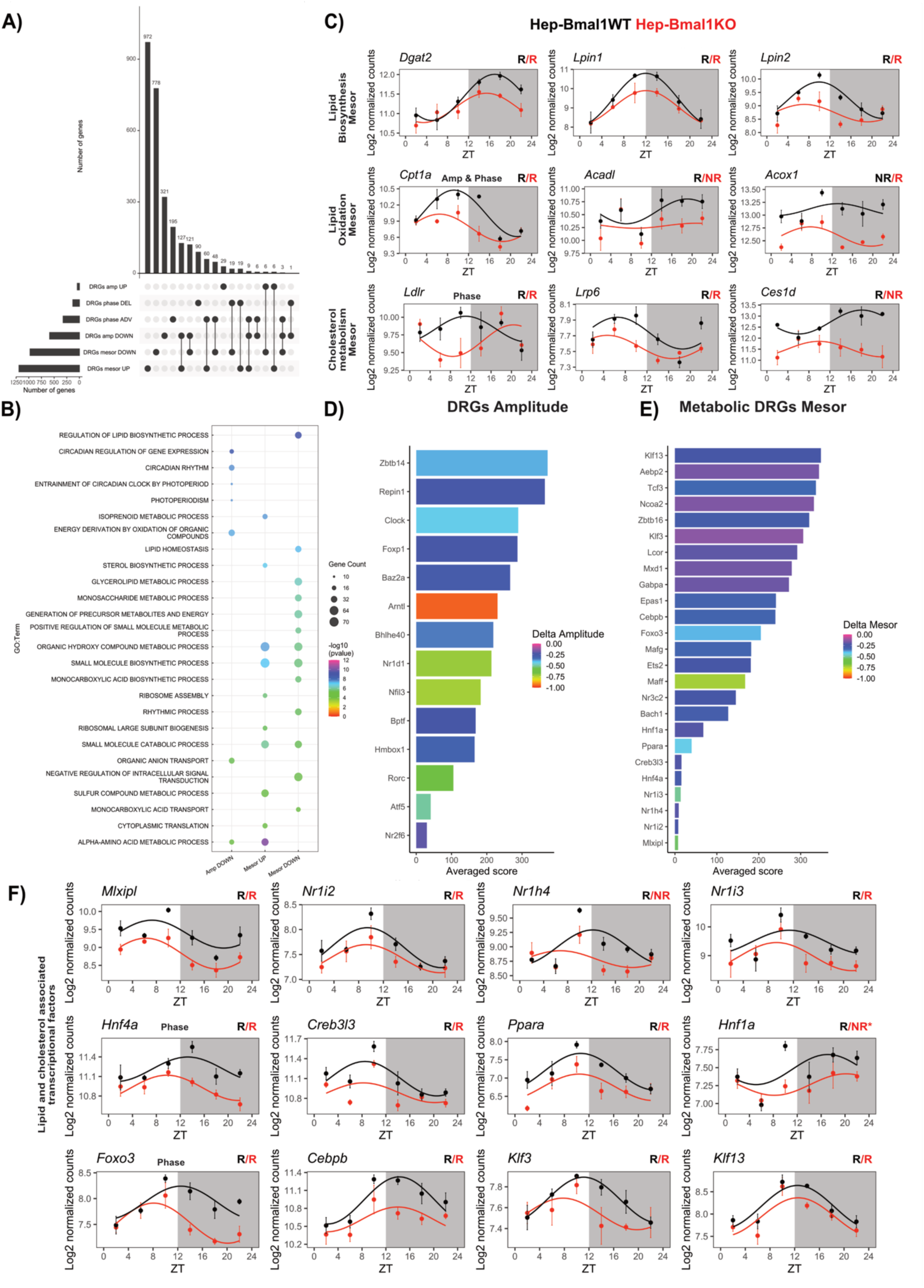
Differential rhythm analysis of the diurnal liver transcriptome in Hep-Bmal1WT and Hep-Bmal1KO mice on a normal chow diet reveals disrupted circadian rhythms and decreased expression of key transcription factors. A) UpSet plot depicts the number of differentially rhythmic genes (DRGs). B) Enrichment analyses of the identified DRGs are depicted. C) Diurnal expression profile of selected DRGs is depicted. D – E) Predictive transcriptional factor analyses using Ch3A3 for DRGs with reduced amplitude or mesor. F) Diurnal expression profile of TFs identified using Ch3A3 algorithm is depicted. Changes in additional rhythmic parameters are described in bold letters. N = 4 per ZT. ‘R’ symbol indicates rhythmicity, while ‘NR’ signifies arrhythmicity detected by CircaCompare.

Since lipid metabolism was strongly ranked in the enrichment analyses, this prompted us to further investigate this pathway. Most of the effects identified in lipid metabolism were associated with reduced mesor, especially in lipid biosynthesis (*Dgat2*, *Lpin1*, *Lpin2*, and *Scd1*) and oxidation (*Cpt1a*, *Acadl*, *Acox1*). Similarly, cholesterol metabolism-associated DRGs (*Ldlr*, *Lrp6*, *Ces1d*, *Ces1e*) showed reduced mesor (Figure 2 C). Considering the role of BMAL1 as a TF, we hypothesized that its loss would affect the rhythmic expression of secondary TFs and impair the expression of lipid- and cholesterol-associated genes. We tested this hypothesis by predicting putative TFs associated with amplitude down DRGs using the ChEA3 algorithm ^16^. By applying stringent criteria (score < 400 & amplitude down), we identified 14 candidate TFs. As expected, *Bmal1* (also known as *Arntl*) was one of these transcription factors exhibiting the greatest reduction in amplitude. We also identified several other clock-associated TFs, including *Clock*, *Bhlhe40*, *Nr1d1*, *Nfil3*, and *Rorc*, which is in line with the pivotal role of BMAL1 in sustaining circadian clock function (Figure 2 D). Subsequently, we focused our analysis on DRGs with reduced mesor that have a direct role in energy metabolism by using the KEGG database. ChEA3 prediction in this subset of genes revealed 25 TFs, among which 12 were identified (e.g., *Chrebp (Mlxipl)*, *Nr1i2*, *Nr1h4*, *Nr1i3*, *Hnf4a*, and *Creb3l3*) as key lipid and cholesterol metabolism regulators, all demonstrating reduced mesor (Figure 2 E – F).

Our findings demonstrate the multifaceted consequences of disrupting the hepatocyte clock in mice fed with normal chow *ad libitum*. Notably, the significant reduction in mesor observed in clock target genes associated with lipid and cholesterol metabolism was associated with expression changes in metabolism regulator TFs.

### Loss of *Bmal1* in hepatocytes worsens MASH

In light of the significant reduction in the expression of several metabolically relevant TFs, we postulated that Hep-Bmal1KO mice would exhibit heightened susceptibility to MASH development. To test this, we subjected Hep-Bmal1KO and -WT mice to a choline-deficient high-fat diet (cdHFD), which was established to induce MASH ^14^. We conducted untargeted lipidomics at the peak time (ZT 6) of endogenous triglyceride and cholesterol levels ^17^. A pronounced diet effect on lipid levels was observed irrespective of genotype. For instance, an increase in cholesterol esters, diglycerides, triglycerides, and phosphatidylglycerols, as well as a reduction in acyl-carnitine, ceramide, lysophosphatidylcholine, phosphatidylcholine, lysophosphatidylethanolamine, and phosphatidylethanolamine species was identified (Figure 3 A, Table S2). Lipids influenced by both diet and genotype included cholesterol esters, plasmalogen phosphatidylcholine, plasmalogen phosphatidylethanolamine, and sphingomyelin species (Figure 3 B, Table S2). Increased cholesterol levels in Hep-Bmal1KO livers were confirmed by an independent method (Figure 3 C). Pathological MASH scoring revealed no marked differences in steatosis and ballooning but higher inflammation scores in Hep-Bmal1KO compared to -WT mice (Figure 3 D-E). Moreover, Hep-Bmal1KO mice fed with cdHFD exhibited a nearly tenfold increase in peri-sinusoidal fibrosis compared to Hep-Bmal1WT controls (Figure 3 F).

**Figure 3:**
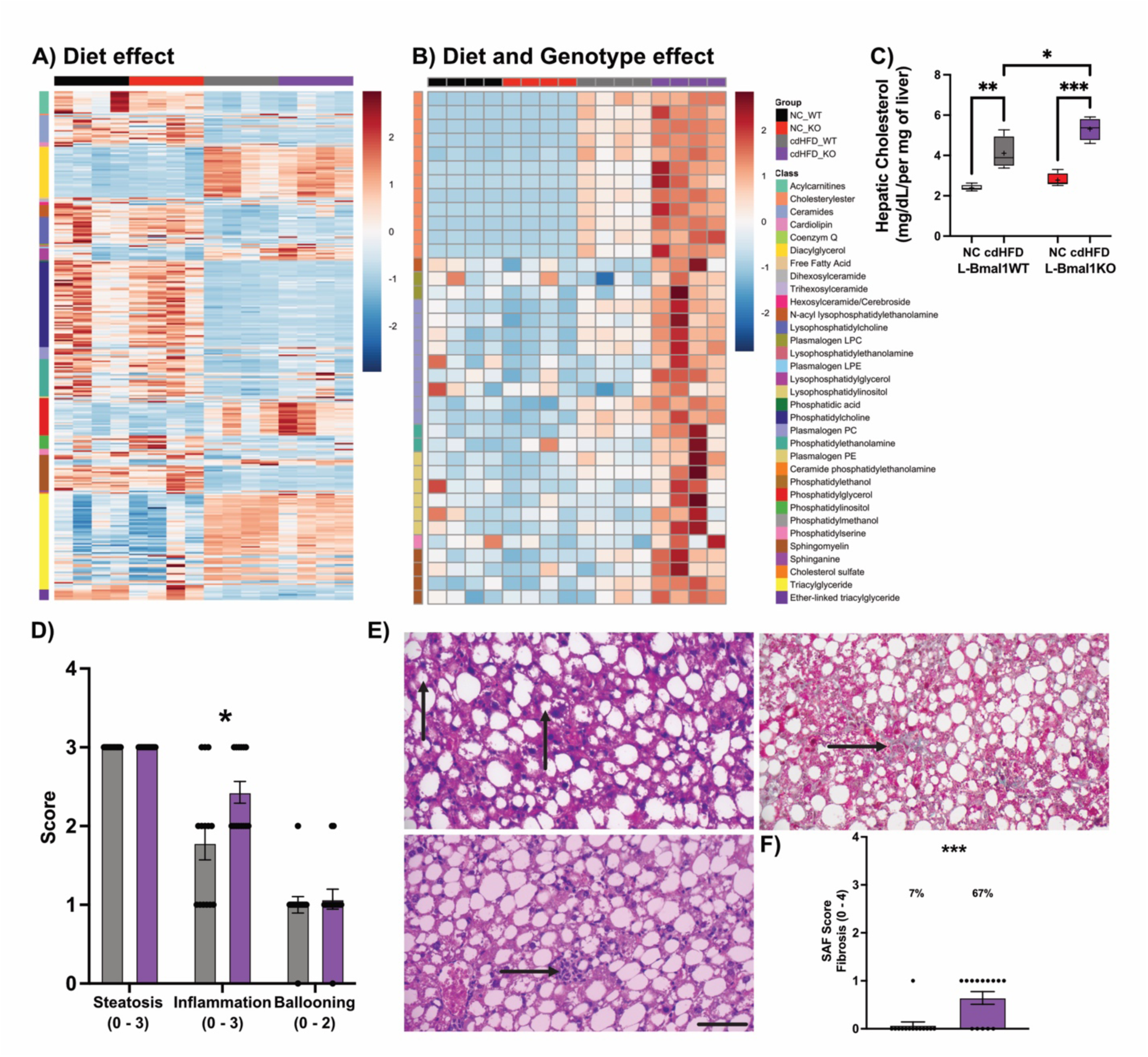
Deletion of *Bmal1* in hepatocytes elevates hepatic cholesterol levels, exacerbating inflammation and fibrosis. A) Heatmap depicts all lipids identified with a diet effect. B) Lipids with a diet and genotype effect are shown by a heatmap. Liver samples were obtained from ZT 6. Normalized lipid species levels and their identity are shown. C) Evaluation of total hepatic cholesterol across genotypes and diets. D) MASLD activity score of Hep-Bmal1WT and Hep-Bmal1KO fed with cdHFD is shown. E) Representative images of ballooning (upper left), inflammation (lower left), fibrosis (upper right), and fibrosis quantification (lower right) are depicted. For histological analysis, samples from the light (ZT 2 - 10) and dark phases (ZT 14 – 22) were used. Scale bar 100 μm.

### Loss of hepatocyte *Bmal1* alters TF gene network rhythms in MASH

To understand how the loss of *Bmal1* in hepatocytes alters cholesterol metabolism and MASH development, we compared daily transcriptional signatures between livers of Hep-Bmal1WT and Hep-Bmal1KO mice under MASH conditions. Rhythm parameter analysis identified several hundred DRGs that exhibited changes in mesor (1,631), amplitude (441), or phase (376) (Figure 4 A). Enrichment analysis of DRGs with decreased mesor revealed pathways related to lipid and fatty acid metabolism, small molecule catabolism, xenobiotic metabolism, and the urea cycle. Conversely, enrichment of DRGs with increased mesor yielded processes such as wound healing, repair, and chondrocyte proliferation. DRGs with reduced amplitude were enriched for cell cycle regulation, DNA replication, cytokinesis regulation, DNA damage response, and circadian rhythms. Processes related to the immune system, including virus response, were overrepresented in DRGs with increased amplitude. DRGs associated with a phase delay were most enriched for circadian processes, while those with phase advancement were enriched for cellular damage response and the Wnt pathway (Figure 4B; Table S3).

**Figure 4:**
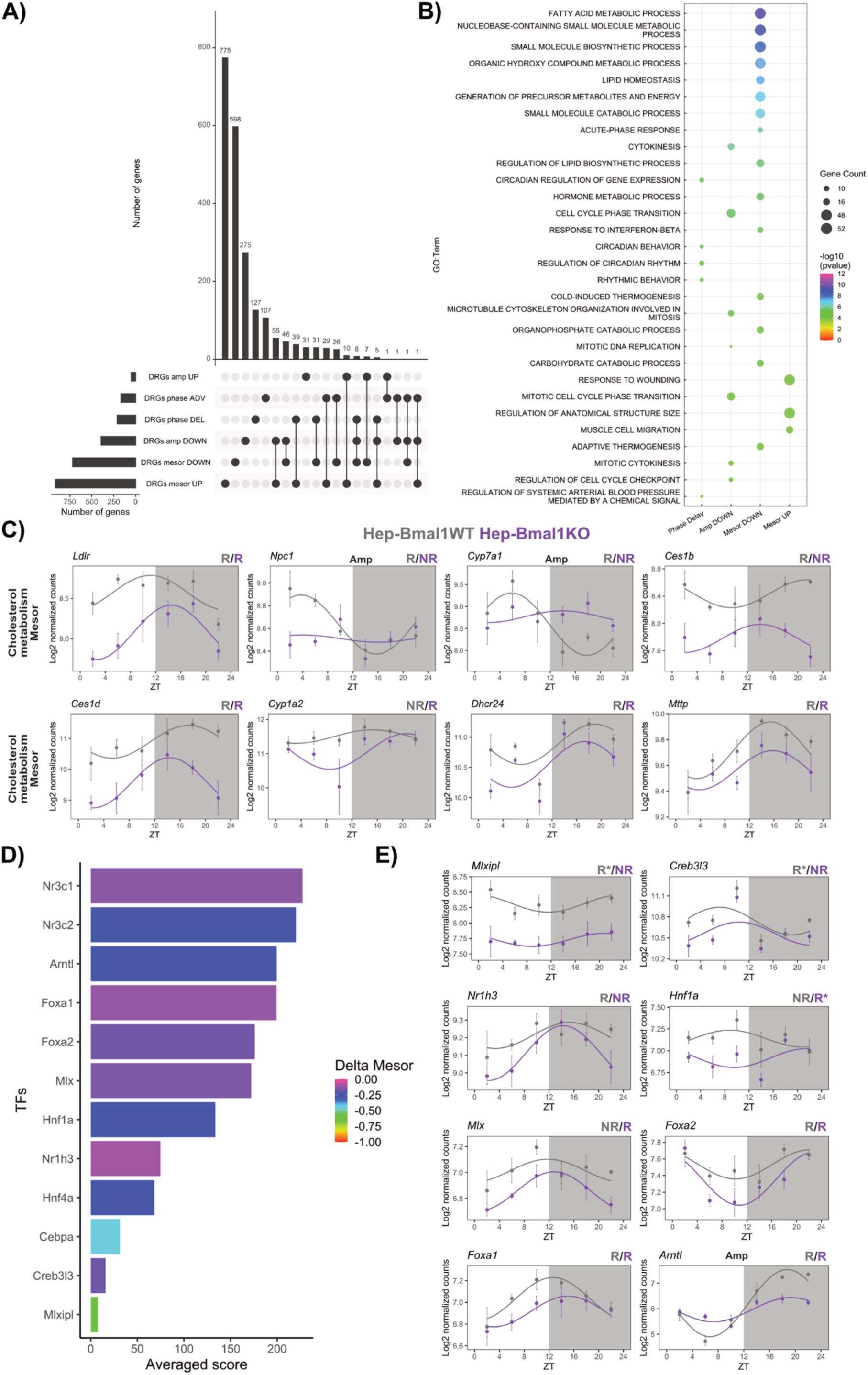
Differential rhythm analysis of the diurnal liver transcriptome in Hep-Bmal1WT and Hep-Bmal1KO mice on a cdHFD shows that *Bmal1* deletion disrupts the expression of cholesterol-related genes. A) UpSet plot depicts the number of differentially rhythmic genes (DRGs). B) Enrichment analyses of the identified DRGs are depicted. C) Diurnal expression profile of selected DRGs associated with cholesterol metabolism. Changes in additional rhythmic parameters are described in bold letters. D) Predictive transcriptional factor analyses using Ch3A3 for DRGs with reduced mesor. E) Diurnal expression profile of transcriptional factors (TFs) identified using the Ch3A3 algorithm is depicted. ‘R’ symbol indicates rhythmicity, while ‘NR’ signifies arrhythmicity detected by CircaCompare.

Based on our lipidomics findings, we performed a targeted analysis of cholesterol metabolism-associated DRGs. Several genes involved in cholesterol uptake (*Ldlr*), intracellular cholesterol transport (*Npc1*), bile acid secretion (*Cyp7a1*), cholesterol metabolism (*Ces1b*, *Ces1d*, *Cyp1a2*, *Dhcr24*, *Mttp*) showed significant reductions in mesor. Additionally, reduced amplitudes of mRNA profiles for *Npc1* and *Cyp7a1* were observed with profiles classified as being non-rhythmic in Hep-Bmal1KO livers (Figure 4 C). We employed the ChEA3 algorithm to identify putative TFs associated with DRGs exhibiting decreased mesor. This analysis was also performed on a subset of energy metabolism-associated DRGs with decreased mesor. We consolidated both lists to focus on shared TFs implicated in metabolism (Figure 4D). A total of 12 TFs were identified (Figure 4 E), among which *Chrebp* (*Mlxipl*), *Creb3l3*, *Cebpa*, *Hnf4a*, and *Nr1h3* were top ranked.

### Hepatocyte clock-driven transcriptional programs regulate cholesterol metabolism

Considering that *Chrebp/Mlxipl* was identified as the top-TF associated with the DRGs exhibiting reduced mesor in our MASH model and its established role in regulating carbohydrate and lipid metabolism ^18^, we hypothesized a cooperative interaction between *Bmal1* and *Chrebp* in regulating MASLD/MASH progression. To directly test this interaction, we used an in vitro setup with murine hepatocytes treated with palmitate to induce steatosis ^19,20^. Following 48-hour palmitate treatment, a transcriptome analysis was conducted to assess the impact of palmitate-induced steatosis on gene expression. 853 genes were up- and 975 downregulated by palmitate (BSA, *vs.* Palmitate, padj < 0.1; Figure 5 A – B; Table S4). Enrichment analysis of the upregulated genes revealed involvement in ribonucleoprotein complex biogenesis, ribosomal RNA processing, protein folding, and post-transcriptional regulation. In contrast, downregulated genes were enriched in pathways related to inflammatory response, cytokine biosynthesis, and lipid, carbohydrate, and steroid metabolism. Notably, palmitate treatment recapitulated several processes observed in Hep-Bmal1WT mice fed a cdHFD, with both conditions exhibiting upregulation of pathways related to cytoskeleton organization, protein transport and folding, and cellular responses to stress. In contrast, processes enriched among downregulated genes in both datasets included pathways involved in lipid metabolism, small molecule biosynthetic processes, and, expectedly, circadian rhythms (Figure 5 C; Tables S3 and S4).

**Figure 5:**
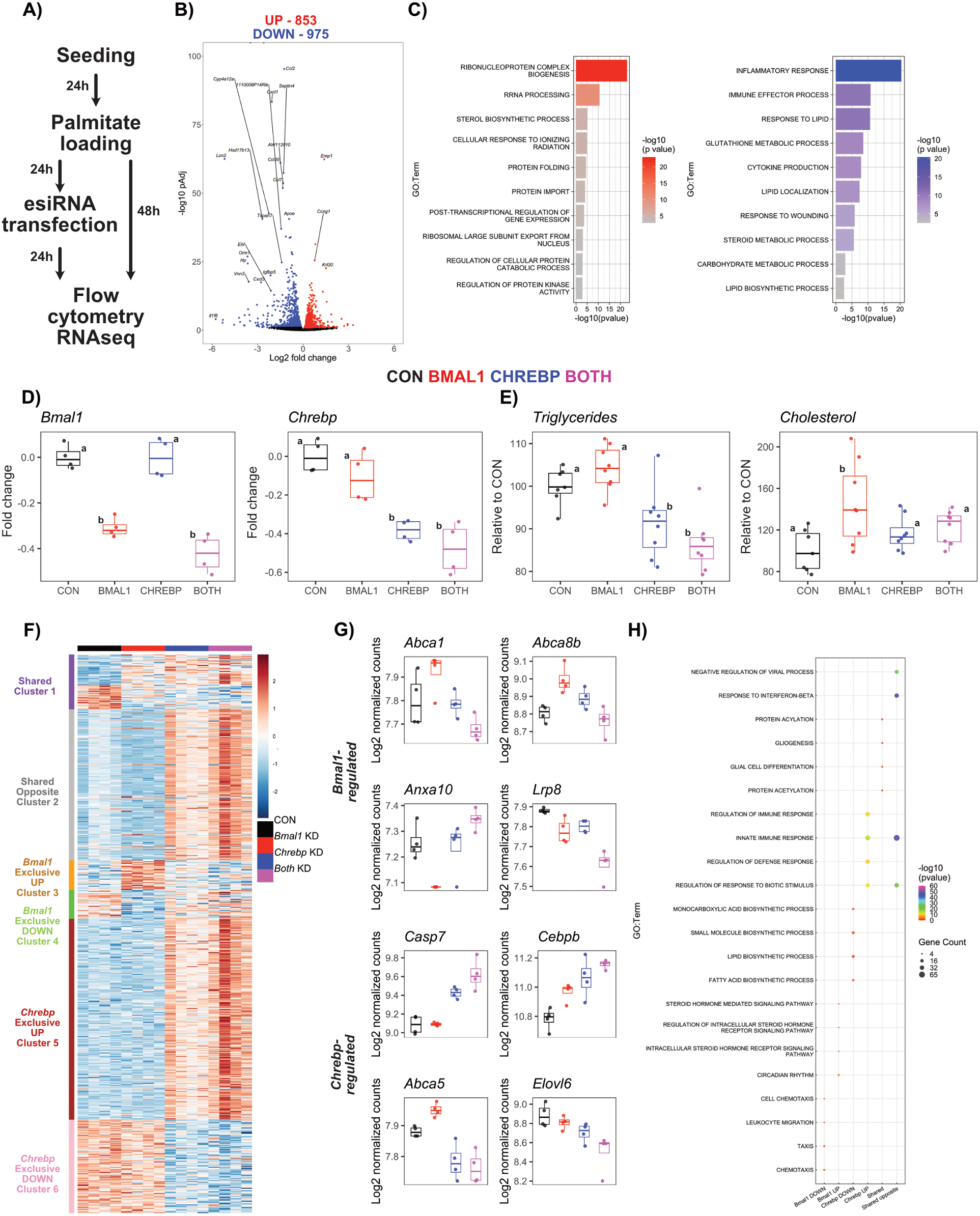
An in vitro model of MASLD demonstrates that the *Bmal1*-regulated transcriptional program specifically controls cholesterol metabolism in immortalized hepatocytes, without impacting lipid metabolism. A) Schematic of the experimental design. B – C) Volcano plots displaying genes with increased (red) and decreased (blue) expression, accompanied by enrichment analysis (n = 4 per group). D) qPCR validation of knockdown efficiency for *Bmal1*, *Chrebp*, and combined *Bmal1*-*Chrebp* (BOTH) (n = 4 per group). E) Dual staining analysis of triglycerides and cholesterol via flow cytometry (n = 7-8 per group). Letters denote statistical significance determined by One-Way ANOVA follow by Tukey post-test. F) Heatmap illustrating gene expression clustering (n = 4 per group), with distinct clusters highlighted in response to gene knockdown (KD). G) Representative plots showcasing unique *Bmal1*- and *Chrebp*-regulated targets. H) Enrichment analysis of the clusters identified in panel F.

To explore a putative interaction between *Bmal1* and *Chrebp*, we conducted combinatory silencing experiments using endoribonuclease-prepared siRNA (esiRNA). Twenty-four hours after esiRNA transfection (48 hours post-palmitate treatment), transcriptome analyses were performed. Transcriptome data confirmed the successful silencing of *Bmal1* and *Chrebp*, which was further validated by qPCR (Figure 5 D; Table S4). DESeq2 analyses identified 1,377 differentially expressed genes (DEGs) in response to *Bmal1*, *Chrebp*, or dual silencing (Likelihood Ratio Test, LRT, padj < 0.1). Dual-labeling triglycerides and cholesterol staining revealed that silencing *Chrebp* alone, or in combination with *Bmal1*, significantly reduced triglyceride levels, indicating a *Chrebp*-driven effect. However, only *Bmal1* silencing resulted in elevated cholesterol levels in murine hepatocytes (Figure 5 E), mirroring the in vivo phenotype observed in cdHFD-fed Hep-Bmal1KO mice (Figure 3 B – C).

To identify genes exclusively regulated by *Bmal1* or *Chrebp*, we filtered the previously identified DEGs using a threshold of a 25%-fold change (log2 of 0.32), ensuring they were unique to either the *Bmal1* or *Chrebp* dataset (Figure 5 F). This analysis yielded 640 DEGs, which were categorized into six distinct classes: shared with similar regulation (63 DEGs), shared with opposite regulation (173 DEGs), genes exclusively upregulated upon *Bmal1* silencing (34 DEGs, Figure 5 G), genes exclusively downregulated upon *Bmal1* silencing (33 DEGs, Figure 5 G), genes exclusively upregulated upon *Chrebp* silencing (230 DEGs, Figure 5 G), and genes exclusively downregulated upon *Chrebp* silencing (107 DEGs, Figure 5 G).

Comparison of the genes co-regulated by *Chrebp* and *Bmal1* in the same direction (shared cluster, Figure 5 F) revealed enrichment of protein acetylation and positive regulation of immune effector process (Figure 5 H; Table S4). In contrast, a distinct set of genes showed opposite regulation when either *Chrebp* or *Bmal1* was silenced (shared opposite cluster, Figure 5 F). These genes were also mainly linked to immune responses, interferon-beta signaling, and cytokine production (Figure 5 H; Table S4). These results demonstrate that both genes significantly influence immune- and inflammation-related gene programs, but in opposing directions. Notably, the number of genes with opposing regulation was roughly three times greater than those with shared regulation.

*Chrebp*-exclusive upregulated DEGs were also associated with immune system responses such as NF-kappaB signaling, regulation of leukocyte differentiation, and cytokine production. On the other hand, *Chrebp*-exclusive downregulated DEGs were primarily associated with fatty acid and lipid metabolism (Figure 5 H; Table S4). Albeit numerically smaller than *Chrebp*-exclusive genes, downregulated *Bmal1*-exclusive genes were enriched in pathways related to innate immune response and chemotaxis, whereas *Bmal1*-exclusive upregulated DEGs were associated with circadian rhythms, steroid metabolism, and lipid metabolism (Figure 5 H; Table S4). Further analysis of *Bmal1*-associated genes revealed several indirect regulators of energy metabolism (e.g., *Igf1*, *Lbh*, *Rorc*, and *Foxa1*) in addition to identifying *Abca1* and *Abca8b*, key cholesterol efflux proteins (Figure 5 F – G).

To further investigate an interaction between BMAL1 and CHREBP, we performed ChipPCR analysis of established *Chrebp* targets (e.g., *Txnip*, *Pklr*, *Klf10*) at the highest time of BMAL1 binding (ZT 6) ^21^. In Hep-Bmal1KO mice, loss of BMAL1 was associated with increased CHREBP occupancy at these promoters, while this pattern was unaffected by the presence of MASH. These findings suggest that hepatic *Bmal1* deficiency enhances CHREBP binding to its target genes, indicating a perturbation of CHREBP-regulated transcriptional programs (Figure S1). Notably, the transcriptomic perturbation, evidenced by the number of DEGs following gene knockdown, was more pronounced in *Chrebp*-silenced samples, suggesting that *Chrebp* has a stronger influence on liver transcriptome regulation. Moreover, we identified several *Chrebp*-regulated gene programs associated with lipid metabolism. The significant downregulation of genes involved in lipid biosynthesis is consistent with the observed reduction in triglyceride levels in *Chrebp*-silenced samples, in line with previous studies on *Chrebp* overexpression or knockdown ^22–24^.

Although *Bmal1* silencing had a relatively modest impact on the overall transcriptome, it notably upregulated several cholesterol-related genes, particularly *Abca1* and *Abca8b*, suggesting a compensatory response to elevated cholesterol levels. This highlights the selective role of *Bmal1* in liver cholesterol metabolism. In vitro data, supported by in vivo diurnal transcriptome analyses, indicate that *Bmal1* deletion disrupts multiple aspects of cholesterol regulation. This includes reduced cholesterol uptake (*Ldlr*, reduced expression), impaired intracellular cholesterol transport (loss of *Npc1* rhythmicity), diminished bile acid secretion (loss of *Cyp7a1* rhythmicity), and increased cholesterol secretion (elevated *Abcg5* and *Abcg8* expression). Taken together, our in vivo and in vitro findings suggest a selective role of *Bmal1* in regulating cholesterol metabolism.

### Patients with MASH show an altered internal circadian phase

In order to translate our findings to the human situation, we evaluated hepatic cholesterol content and circadian clock gene expression in liver biopsies from a patient cohort living with obesity, and either metabolic dysfunction-associated steatotic liver (MASL, simple steatosis) or MASH with a body mass index (BMI) ≥ 40 or ≥ 35 kg/m^2^ with at least one cardiometabolic comorbidity (hypertension, type 2 diabetes mellitus (T2DM), dyslipidemia or obstructive sleep apnea syndrome (OSAS) (Table S5)). To account for temporal effects on clock gene expression, we developed an internal phase index by multiplying *PER3* expression with *NR1D2*, then dividing the result by *BMAL1* expression. A higher ratio implies an increased expression of *PER3* and *NR1D2*, whereas a decreased ratio suggests increased *BMAL1* expression (Figure S2). We identified that patients with MASH had a higher ratio compared to patients with MASL (e.g., simple steatosis), thus suggesting altered internal phasing in MASH (Figure 6 A). Importantly, both MASL and MASH samples were collected across the day and not restricted to a given phase, thus reducing a temporal bias in our sample collection. However, no differences in total hepatic cholesterol between MASH and MASL samples were observed (Figure 6 A). Considering the entire cohort (MASL and MASH), hepatic cholesterol was positively correlated with serum HDL levels. Internal phasing had a borderline significant trend with disease score (NAS, p = 0.07, Figure 6 B).

**Figure 6:**
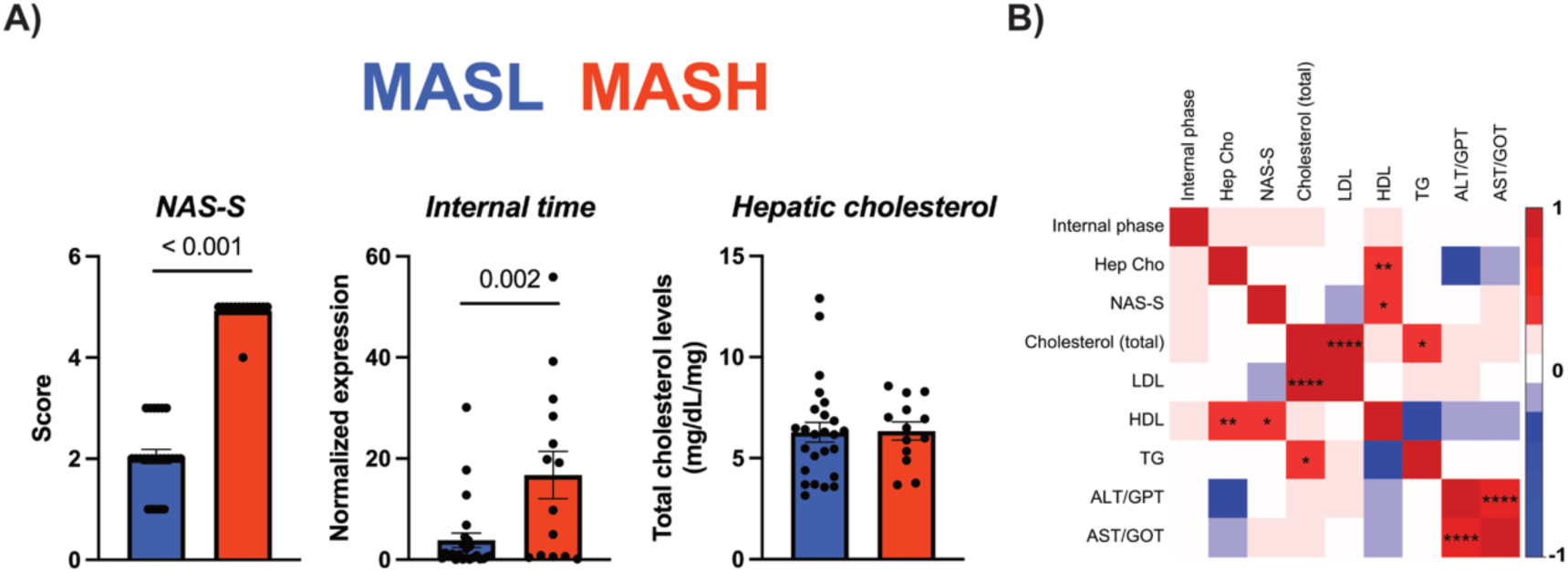
Altered Circadian Phase in Livers of MASH Patients Compared to Simple Steatosis. A) NAS-Score, internal timing expression, and hepatic cholesterol levels of MASL and MASH patients are depicted. B) Correlation of clinical data with hepatic cholesterol and internal timing. N = 14 for MASL and 24 for MASH.

Therefore, the human findings confirm a correlation between disrupted circadian rhythms, namely altered internal phasing, with MASL-to-MASH progression.

## DISCUSSION

Our research highlights the role of the hepatocyte circadian clock. Under physiological conditions, the absence of the circadian clock in hepatocytes led to a reorganization of the hepatic transcriptome, impacting thousands of genes with alterations in mesor, phase and/or amplitude. It further decreased the expression of several metabolic-associated TFs, suggesting a possible interaction between BMAL1 and other TFs. The protective function of the hepatocyte clock became apparent when mice were subjected to cdHFD, as Hep-Bmal1KO mice demonstrated exacerbated MASH development. At the transcriptional level, the absence of the hepatocyte circadian clock significantly modified cholesterol metabolism, influencing several genes implicated in cholesterol uptake, transport, and degradation. Predictive bioinformatic analyses identified a potential interaction between *Chrebp* and *Bmal1*. Subsequent in vitro mechanistic studies revealed that *Bmal1* and *Chrebp* share an antagonistic relationship, with these TFs modulating multiple metabolic pathways in opposite directions. A novel aspect of our findings revealed that the hepatic *Bmal1*, under our experimental conditions, selectively regulates cholesterol metabolism. Importantly, our investigation in a cohort of obese MASL/MASH patients had an altered internal phasing, suggesting disrupted circadian rhythms in MASH livers, but did not recapitulate the hepatic cholesterol phenotype observed in experimental models.

Under normal chow conditions, the livers of Hep-Bmal1KO mice exhibited significant disruption in several clock genes, including *Bmal1*, *Clock*, *Cry1*, *Cry2*, *Nr1d1*, and *Nr1d2*. However, it is noteworthy that approximately 50% of the transcriptome remained rhythmic in both Hep-Bmal1KO and Hep-Bmal1WT mice. This observation must be reconciled with the fact that Hep-Bmal1KO mice display rhythmic locomotor and feeding behaviors ^15^. Given that the liver’s circadian clock is particularly sensitive to feeding regimens ^25^, the presence of such consistent rhythms is not unexpected. Comprehensive circadian analyses revealed a pronounced rewiring effect. Among the numerous identified processes, the absence of *Bmal1* in hepatocytes resulted in diminished expression of genes linked to lipid and glycerolipid metabolism. Additionally, phase effects, characterized by advances or delays, were noted in lipid and cholesterol metabolism, suggesting a complex regulation of these pathways. However, a minor effect on the lipidome was observed between Hep-Bmal1WT and Hep-Bmal1KO fed with *ad libitum* normal chow. A notable finding, however, was the substantial downregulation of several TFs. While rhythmicity prevailed in most instances, several TFs exhibited a marked reduction in mesor, indicating *Bmal1’s* regulatory role in the diurnal regulation of these TFs.

Extensive research on the effects of *Chrebp* regulating energy metabolism has been performed. Global *Chrebp* knockout increased plasma glucose due to reduced insulin sensitivity, increased hepatic glycogen levels, and reduced lipogenesis, events which were enhanced when mice were fed a high-starch diet ^26^. In the leptin-deficient model (ob/ob), global *Chrebp* knockout decreased hepatic fatty acid synthesis, normalized triglyceride levels, and reduced body weight^27^. Similar findings were observed in ob/ob mice with hepatocyte-targeted adenovirus-silenced *Chrebp*, including a reduction in hepatic cholesterol levels ^28^. Global *Chrebp* knockout mice fed with a Western diet had reduced hepatic lipid and cholesterol levels, decreased beta-oxidation and ketogenesis, and increased intestinal lipid absorption ^24^. Lipid accumulation in hepatocyte-specific *Chrebp* knockout mice was similar to that in control mice when they consumed regular chow or a high-fat diet. However, these knockout mice exhibited lower lipid levels when fed a carbohydrate-rich diet. Despite being protected from carbohydrate-induced hepatic steatosis, the hepatocyte-specific *Chrebp* knockout mice showed impaired glucose tolerance ^29^. Conversely, hepatic *Chrebp*-overexpressing mice fed a high-fat diet had more pronounced hepatic steatosis despite having improved insulin signaling and glucose tolerance compared to controls ^22^.

The putative interaction between *Bmal1* and *Chrebp* has not yet been explored in depth. Predictive transcription factors linked to genes exhibiting reduced mesor under both control and cdHFD conditions have identified *Chrebp* as the most significantly associated TFs factor in the livers of Hep-Bmal1KO mice. Our in vitro model of steatosis utilizing hepatocytes further illuminated this novel interaction. Transcriptome analysis indicated that *Chrebp* and *Bmal1* likely interact through opposing regulatory pathways, which is evident from the fact that DEGs with opposing regulation are approximately three times higher than those with shared regulation. Silencing *Chrebp* in vitro resulted in a greater disruption, evidenced by nearly a five-fold increase in DEGs compared to *Bmal1*-silenced hepatocytes. Consistent with earlier research ^22,26–28^, *Chrebp* silencing diminished genes related to lipogenesis. Further confirmatory findings came from the in vitro palmitate model in which *Chrebp*-silenced hepatocytes had reduced triglyceride levels compared to control and *Bmal1*-silenced hepatocytes. Importantly, dual *Bmal1* and *Chrebp*-silencing also resulted in reduced triglycerides, thus suggesting a *Chrebp*-dependent effect in line with the transcriptome predictions. Furthermore, the findings supporting the *Chrebp*-*Bmal1* interaction revealed that CHREBP-chipPCR showed enhanced binding of CHREBP to classical CHREBP-regulated genes (*Txnip*, *Pklr*, *Klf10*) in Hep-Bma1KO livers under normal chow conditions, which remained unchanged by MASH. Therefore, our findings identified a novel interaction of *Chrebp* with *Bmal1*, thus suggesting that *Chrebp* effects in the liver can be time-of-day dependent, a matter for future investigation.

Our research highlights the protective role of the hepatic clock during MASH development. Interestingly, at the transcriptional level, many lipid-associated genes had reduced mesor and amplitude in Hep-Bmal1KO mice, but these changes did not affect lipid levels across the genotypes. In contrast, cholesterol-associated genes were affected, and hepatic cholesterol levels were higher in Hep-Bmal1KO mice. Poorer MASH scores and increased fibrosis were also noted in Hep-Bmal1KO mice. Our findings indicate a role for *Bmal1* in regulating cholesterol metabolism, supported by in vitro experiments with *Bmal1*-silenced hepatocytes. The selective role of *Bmal1* in regulating cholesterol metabolism is unclear from previous studies. For example, Hep-Bmal1WT and Hep-Bmal1KO fed with a chow diet exhibited decreased triglyceride levels only in the morning (ZT 0), and no cholesterol levels were measured ^30^. In our experiments (ZT 6), no changes in lipids were noted, except for a few types of glycerophospholipids. A recent study revealed that the silencing of *Bmal1* in hepatocytes enhances m6A mRNA methylation, contributing to the lipid disruption seen in Hep-Bmal1KO mice ^31^. In an elegant study, global *Bmal1* KO mice in an *Apoe*-null background exhibited elevated hepatic cholesterol and triglyceride levels and reduced cholesterol excretion in bile and feces. In Hep-Bmal1KO/ApoeKO mice, hepatic cholesterol and triglyceride levels were also elevated compared to Hep-Bmal1WT/ApoeKO mice. Overexpression of *Bmal1* in the liver of Hep-Bmal1KO/ApoeKO mice increased cholesterol and bile acid levels in the bile. This effect was dependent on *Bmal1* regulating the rhythmic expression of *Abcg5* and *Abcg8* ^32^. Notably, our findings support earlier research by Pan et al. (2016) and reinforce *Bmal1’*s involvement in cholesterol regulation. However, Pan’s study was conducted in an *Apoe*-null background, whereas our research concentrated on the role of *Bmal1* in hepatocytes. Some studies have suggested potential pathways for the protective role of the hepatic clock. For instance, a study has shown that the protective effects of the hepatocyte clock are mediated via TGF-β signaling that suppresses fibrotic gene expression in healthy conditions. Metabolic stress disrupts this rhythmic control, enhancing profibrotic pathways and promoting liver fibrosis in MASH ^33^. Conversely, disruption of the hepatic clock has been shown to disrupt mitochondrial structure, leading to increased oxidative stress, which contributes to insulin resistance and metabolic dysfunction ^34^. Using our in vitro steatosis model, we could identify that hepatocytes treated with palmitate, a mimic for steatotic condition ^19,20^, had increased cholesterol levels when *Bmal1* was silenced compared to control and *Chrebp*-silenced groups. Transcriptome analysis also identified an upregulation of cholesterol efflux transporters (*Abca1* and *Abca8b*), suggesting a compensatory mechanism due to the high cholesterol levels in vitro. Interestingly, the gene signature associated with cholesterol in vivo was associated with reduced mesor and/or amplitude for cholesterol uptake, intracellular transport, bile acid secretion, and higher cholesterol secretion. Collectively, both in vitro and in vivo evidence point towards a selective role of *Bmal1* in regulating cholesterol metabolism.

Circadian rhythms evaluation in MASLD patients is challenging due to the difficulty of obtaining liver samples from the same individual at multiple time points throughout the day. However, in an elegant study by Johanns and colleagues, the authors successfully collected liver biopsies from healthy, MASL, and MASH patients throughout the day. Their analysis revealed time-of-day-dependent transcriptomic changes between morning (AM) and afternoon (PM) samples, regardless of disease status. They also observed that the AM-to-PM fold changes of common time-dependent DEGs were progressively attenuated in steatotic livers and further diminished in MASH livers, compared to healthy controls ^35^. This is in line with the predictions of a gradual disruption of hepatic rhythms as MASLD progresses ^6^. Our findings in humans further support these results, as patients with MASH exhibit altered internal phasing compared to those with MASL, which suggests a further impairment of liver circadian rhythms in MASH compared to MASL.

Our study has its limitations. For instance, our differential rhythm analysis was based on an uncorrected p-value, which may result in higher false positive rates. To address this, a more stringent p-value threshold (<0.01) was set to determine rhythmicity, with these genes later used for differential rhythm analysis. Our analyses did not include the zonation effect. Triacylglycerols, diacylglycerols, sphingolipids, and ceramides exhibited specific alterations and a shift from pericentral to periportal localization in MASH ^36^. However, our earlier research did not find a distinct zonation effect regarding lipid accumulation in mice fed a cdHFD ^14^. This phenomenon is related to the strong induction of MASH by cdHFD, which can occur in just 2 weeks. However, it does not accurately reflect the progression of human MASH, limiting the clinical applicability of our results ^37,38^. In fact, the hepatic cholesterol phenotype induced by *Bmal1* loss was not replicated in the human cohort. This difference could be due to various factors, including differences in hepatic zonation, as biopsy samples likely vary in their representation of metabolic regions with distinct lipid profiles. The analysis of total rather than specific cholesterol fractions, and the early disease stage with minimal fibrosis in all patients, may have precluded detectable cholesterol accumulation at the time of sampling. We could not make a full comparison because there were no healthy lean liver biopsy biopsies available, owing to ethical limitations. Although our target *Bmal1* deletion in hepatocytes led to marked effects in the liver, the contribution of other non-hepatocyte cells in regulating cholesterol metabolism remains elusive and should be addressed using single-cell RNAseq in future experiments.

Taken altogether, our study offers a valuable overview of the role of the hepatocyte clock under normal chow and MASH conditions, providing a useful resource for future experiments. We discovered that impairing the circadian clock of hepatocytes impacts the expression of various metabolism-associated TFs, contributing to the exacerbated development and progression of MASH. We also identified a new connection between *Bmal1* and *Chrebp*, which, while not directly affecting cholesterol metabolism, highlighted a potential circadian regulation of *Chrebp* in the liver. Importantly, these findings are linked to a specific role of *Bmal1* in regulating cholesterol metabolism. Our experimental findings were supported by clinical data showing that human MASH livers show altered internal phasing compared to MASL livers. Collectively, our findings suggest that targeting the circadian clock can be a promising pathway to treat MASH in humans.

## MATERIAL AND METHODS

### Mouse model and experimental conditions

To achieve hepatocyte-specific deletion of *Bmal1*, we used Alb^Cre^ mice (B6.Cg-Speer6-ps1Tg(Alb-cre)21Mgn/J, Jackson Laboratory, stock #003574) crossed with Bmal1^flx/flx^ mice (B6.129S4(Cg)-Bmal1-tm1Weit/J, Jackson Laboratory, stock #007668), as initially described^15^. This breeding generated hepatocyte-specific *Bmal1* knockout mice (Alb^Cre/+^/Bmal1^flx/flx^) and their littermate controls (Alb^+/+^/Bmal1^flx/flx^) referred to as Hep-Bmal1KO and Hep-Bmal1WT, respectively. Genotyping was carried out using specific primers recommended by the Jackson Laboratory. Eight-to-ten weeks old mice were housed under a 12-hour light/12-hour dark cycle at 22 ± 2°C, with free access to food and water, and acclimated to the experimental conditions for at least one week. Mice were fed either normal chow (NC, 27% protein, 14% fat, 59% carbohydrate, Altromin, Germany) or a choline-deficient high-fat diet (cdHFD, 9 % protein, 60 % fat, 2 % cholesterol, and 31 % carbohydrate, with 0.17 % methionine, Ssniff E15673-94, Germany) for two weeks to induce MASH, as previously described ^14^. After the diet period, mice were euthanized every four hours for tissue collection, starting two hours after lights on (ZT 2), which was immediately flash-frozen in liquid nitrogen and stored at -80°C. The study was ethically approved by the Animal Health and Care Committee of the Government of Schleswig–Holstein, following international animal welfare guidelines. Both male and female mice were used in balanced cohorts for the experiments, except group Hep-Bmal1WT fed with cdHFD at ZT14, where it had one female sample. Data reporting follows ARRIVE guidelines.

### Human Cohort

The study enrolled patients from two different medical centers, the Department of General, Visceral, Thoracic, Transplantation, and Pediatric Surgery, University Medical Center Schleswig-Holstein, Kiel and the Department of Surgery, University Medical Centre Schleswig-Holstein, Campus Lübeck. All patients with obesity had undergone surgical liver biopsy during the bariatric intervention. The patients were aged >18 years and fit the criteria for obesity surgery according to current guidelines, namely, body mass index (BMI) ≥ 40 or ≥ 35 kg/m^2^ and at least one cardiometabolic comorbidity (hypertension, T2DM, dyslipidemia or OSAS). Pre-operative evaluation included a detailed medical history, physical examination and nutritional, metabolic, cardiorespiratory and psychological assessment. Exclusion criteria were autoimmune, inflammatory or infectious diseases, viral hepatitis, cancer or known alcohol consumption (>20 g/day for women and >30 g/day for men). Groups were categorized according to liver histology. Clinical data and type of bariatric surgery are summarized in Table S5. All patients provided written informed consent. The local ethics committee approved the present clinical investigations. The study was conducted in accordance with the Declaration of Helsinki. Liver samples were either snap-frozen directly in liquid nitrogen in the operating theatre or stored in PBS on ice for a short period for transport before freezing. Samples were stored at -80 °C until further processing.

### Bulk Transcriptome analyses

Transcriptome analysis was performed using the Bulk RNA Barcoding and sequencing (BRB-seq). In this method, only the 3’ portion of the mRNA is sequenced, thus reducing the need to sequence the whole transcript, and it has a similar efficiency compared to TruSeq method ^39^. RNA from livers was extracted using Trizol according to the manufacturer’s recommendation. High-quality RNA with ratios (260/280 and 260/230 ratios) higher than 1.8 were used. RNA integrity was assessed by gel electrophoresis. Similar RNA input was used to generate cDNA libraries using the MERCURIUS kit (Alithea Genomics). Libraries were sequenced on Illumina NovaSeq 6000 platform at a depth of 8 million raw reads per sample. The sequencing reads were demultiplexed using the BRB-seq tools suite and aligned against mouse genome (mm10) using STAR and count matrices were generated using HTSeq ^39^. FeatureCounts was used to count the read numbers mapped to each gene.

### Lipidomics analysis of the liver via liquid chromatography coupled to tandem mass spectrometry (LC-MS/MS)

Liver tissue was cryogrinded using a Cellcrusher device. 50µl of water was added to 20 mg of tissue powder. Lipidomics analyses were performed as previously described ^19^. In brief, measurements were conducted with a Dionex Ultimate 3000 RS LC-system and an Orbitrap mass spectrometer, using high-quality solvents and a heated-electrospray ionization probe. Lipid profiling involved extracting lipids from tissue homogenate using a specific solvent mix, followed by LC-MS/MS analysis on an Accucore C30 RP column. Data was acquired with data-dependent MS² scans and lipids were identified using Compound Discoverer 3.3 and two additional databases ^40^. The area under the peak was normalized to the internal standard and sample weight. An extraction blank and quality control samples were used to ensure data quality and linearity. Raw data were log2 transformed and analyzed using two-way ANOVA followed by Tukey’s Honest Significant Difference (HSD), corrected by multiple comparisons. Validation of cholesterol levels was performed using a commercial kit (STA 384; Cell Biolabs). Two-way ANOVA followed by Tukey’s post-test was used to analyze the data. In both cases, padj < 0.05 was deemed significant.

### Hepatic cholesterol evaluation

Total hepatic cholesterol was processed according to the manufacturer’s instructions (Cell Biolabs, San Diego, USA, STA 384). In short, liver samples (approximately 10 mg) were homogenized in 0.2 mL of extraction buffer (chloroform:isopropanol:NP40, 7:11:0.1) using a bead-based homogenizer. Following centrifugation at 15,000 × g for 10 minutes at room temperature, the supernatant was carefully transferred to new tubes, ensuring avoidance of the pellet. Samples were incubated at 50°C for 30 to 60 minutes and vacuum-dried at 45°C for 30 to 60 minutes. Dried lipids were resuspended in 200 μL of 1× Assay Diluent. The Cholesterol Reaction Reagent was freshly prepared by combining Cholesterol Oxidase (1:50), HRP (1:50), Colorimetric Probe (1:50), and Cholesterol Esterase (1:250) in 1× Assay Diluent. Cholesterol standards were created by serial dilution of a 250 μM cholesterol stock solution in 1× Assay Diluent. Fifty microliters of samples diluted 1:20 were added to a 96-well plate, followed by 50 μL of Cholesterol Reaction Reagent. After incubation at 37 °C for 45 minutes in the dark, absorbance was measured at 555 nm, and cholesterol levels were determined using a standard curve. Hepatic cholesterol was normalized by the amount of tissue. Permutation test using the package lmPerm in R studio was used to compare human data.

### Histological analyzes

For H&E staining, mouse liver samples were collected, rinsed in PBS, and fixed in 4% paraformaldehyde (PFA, Electron Microscopy Sciences) for 48 hours. Subsequently, the samples were dehydrated through a graded ethanol series. The livers were then embedded in paraffin and sectioned into 2-5 µm slices. Following dewaxing and rehydration, the slices were stained with Hematoxylin and Eosin (H&E) for examination under a light microscope. Elastica van Gieson (EvG) Staining and Gömöri Silver Impregnation was used. A senior pathologist, who was blinded to the sample identities, assessed the sections and assigned scores using the NAFLD Activity Score (NAS) and the Steatosis, Activity, and Fibrosis (SAF) scoring systems. A Mann-Whitney test was used to compare the groups.

### Rhythm analyses

Genes containing a sum of reads among the groups lower than 100 were excluded from the analysis. PCA method was used to assess possible outliers. No sample exclusion was performed, and all 96 samples were considered for the downstream analysis. The remaining genes were log2 transformed in DESeq2 using the vst function ^41^. A total of 15,548 transcripts were considered for subsequent analyses. To analyze the diurnal patterns in gene expression, we employed four distinct algorithms: CircaN ^42^, Jonckheere-Terpstra-Kendall (JTK) cycle ^43^, Metacycle ^44^, and DryR ^45^. CircaCompare was utilized for the comparative analysis of rhythmic parameters across different groups ^46^. For the evaluation of mesor and amplitude, CircaCompare was applied to fit a sine curve without the necessity for rhythmicity criteria fulfillment. Phase comparisons were specifically conducted for genes exhibiting robust rhythmicity. The algorithms CircaN, JTK cycle, and Metacycle were run within a CircaN framework, using standard settings and assuming a fixed 24-hour period. We established a threshold for rhythmicity significance at p < 0.01 for each analytical method. Subsequently, rhythmic genes identified in at least one of these methods were aggregated into a unified list for each group, and rhythm parameters were compared utilizing the CircaCompare algorithm using a p < 0.05.

### Enrichment analysis

Gene lists were analyzed using the ClusterProfiler package in R environment, with the entire identified genes serving as the background dataset. For cases where identified biological processes exceeded 100 hits, the simplify function was applied with a cutoff value of 0.6 to streamline the results. In scenarios with fewer processes, only those involving more than three genes and exhibiting a p value < 0.01 were considered for further analysis.

### ChIP-X Enrichment Analysis Version 3 (ChEA3)

TFs prediction was performed using the online application ChEA3 (https://maayanlab.cloud/chea3/) ^16^ using DRGs as input. Metabolic DRGs were selected using the KEGG term “Metabolic Pathways”. Mean library rank was used. TFs were filtered for a score < 400.

### In vitro experiments

AML-12 cells, obtained from ATCC Biobank (catalog no. CRL-2254), were cultured in Gibco Dulbecco’s Modified Eagle Medium supplemented with Nutrient Mixture F12 (DMEM/F12, ThermoFisher), containing 1% penicillin-streptomycin, 1% insulin-transferrin-selenium (ITS, ThermoFisher), 10% non-heat-inactivated fetal bovine serum (FBS, ThermoFisher), and 10 nM dexamethasone (Sigma-Aldrich). Cells were incubated at 37 °C in a humidified atmosphere with 5% CO2. To mimic a steatosis condition, cells were cultivated in media without dexamethasone and FBS was replaced by charcoal/dextrane-treated FBS (Hyclone). One hundred thousand cells were seeded per well in 12-well plates and 24 hours later received palmitate-BSA (0.25 mM, 7:1 Cayman Chemical, USA). On the following day, cells were transfected with esiRNA against *Bmal1* (100 nM), *Chrebp* (100 nM), or both genes (200 nM) in addition to the negative control esiRNA (200 nM *Egfp*, Euphoria Biotech, Germany) using lipofectamine 3000 according to the manufacturer’s recommendation (Thermofisher, USA). Twenty-four hours after esiRNA addition, cells were collected and processed for transcriptome analysis (BRB-seq – see above) or flow cytometry, as described next.

Dual triglyceride and cholesterol labeling was performed as described before ^19^. In brief, cells were detached from the wells using Trypsin-EDTA (ThermoFisher, USA), pelleted by centrifugation (500 x g per 5 min) and incubated sequentially with Zombie NIR dye (1:500 dilution, Biolegend, USA), Bodipy-Cholesterol (1 μM, MedChem Express, Sweden), and AdipoRed (0.3 μL per mL, Lonza, Switzerland) at room temperature. Each incubation step was combined with centrifugation (1000 x g per 3 min) and washing with PBS. Cells were then fixed in 4% PFA for 30 minutes at room temperature and stored for flow cytometry analysis. Samples were assessed in Cytek Aurora at the Cell Analysis Core Facility (CAnaCore facility at the University of Lübeck). Data were acquired using predefined templates with adjustments for optimal event rates and detector gains. Spectral unmixing was applied using the manufacturer’s software with single-stained controls to resolve fluorescent populations. At least 10,000 events were captured for each sample. Gating strategies were implemented using FlowJO V10 software, with each gate clearly defined in terms of fluorescence.

### Differentially expressed genes (DEGs) and clustering strategy

DESeq2 ^41^ was used to analyze differentially expressed genes (DEGs) based on raw count data. Genes with counts lower than 25 were excluded. Pairwise comparisons, such as BSA vs. PAL, were performed using the Wald test with standard parameters (padj < 0.1). To identify genes regulated by *Bmal1* and *Chrebp*, the likelihood ratio test (LRT) was conducted in DESeq2. Unsupervised clustering was then applied using the complete method and Euclidean distance method. *Bmal1*- and *Chrebp* -specific target genes were further refined by applying a log2 fold change threshold (0.32) and ensuring that each gene was exclusively expressed in only one group. The resulting clusters were then subjected to functional enrichment analysis using clusterProfiler.

### Quantitative (q)PCR

RNA purity was confirmed by obtaining 280/260 and 260/230 absorbance ratios greater than 1.8 using a spectrophotometer. Up to 2 μg of the total RNA was reverse transcribed with random hexamer primers using the High-Capacity cDNA Reverse Transcription Kit (Thermo Fisher Scientific, USA). qPCR was performed using the Go Taq qPCR Master Mix (Promega, USA) using 50 ng of cDNA. The following amplification program was used on a Bio-Rad CFX96 cycler (Bio-Rad, Hercules, CA, USA): 5 min at 94◦C, 45 cycles of 15 at 94◦C,15s at 60◦C, and 20s at 72◦C, and final extension for 5 min at 72 ◦C. After amplification, a melt curve was generated to verify product specificity by heating the product from 65◦C to 95◦C at 0.5◦C/s. For human gene expression, the TaqMan method with specific genes and probes (IDT, USA) was employed. The qPCR conditions included an initial activation step of 5 minutes at 95°C, followed by 15 seconds at 95°C, and then 45 cycles of 60 seconds at 59°C and 30 seconds at 72°C. All genes and probes are detailed in Table 1. Relative expression ratios for each transcript were calculated based on individual primer efficiencies using the Pfaffl method ^47^.

**Table 1:**
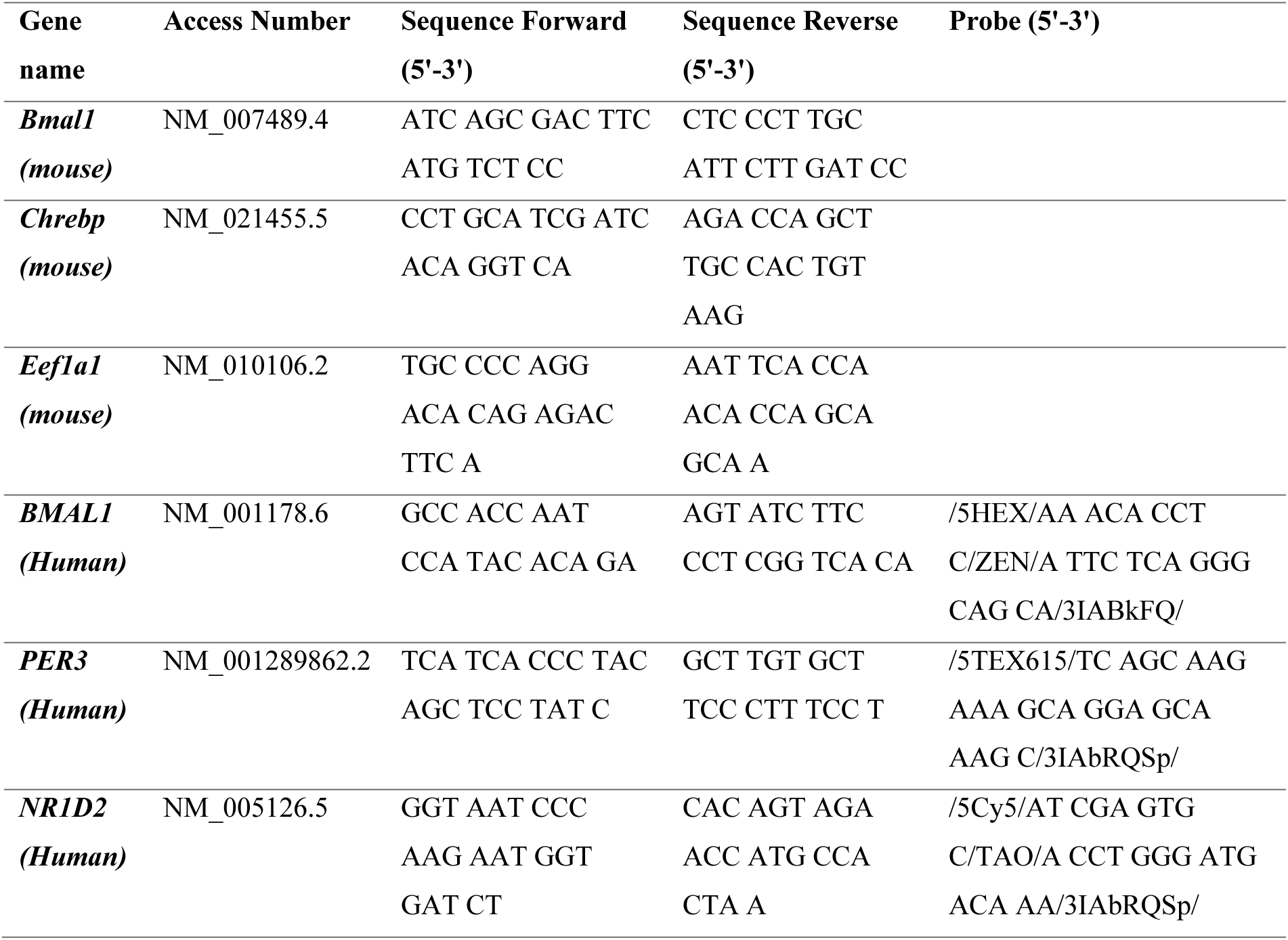
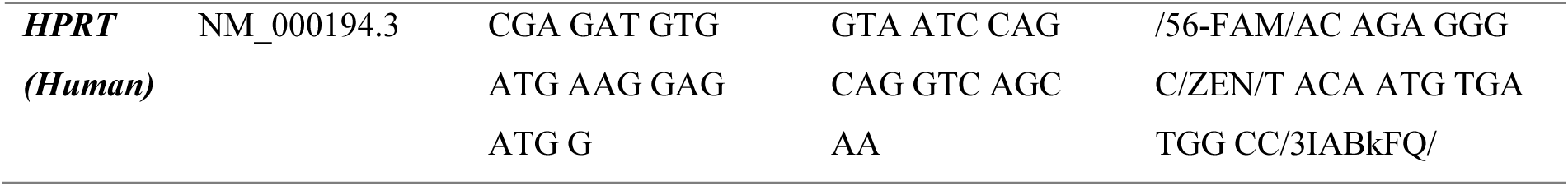
Primers and probes for qPCR.

### ChREBP ChIP-PCR

ChIP was performed as described previously ^48^. In short, small pieces of frozen liver tissue were pulverized in liquid nitrogen and underwent cross-linking in 1% formaldehyde for 10 min, followed by quenching with 1/20 volume of 2.5 M glycine solution. Nuclear extracts were prepared by homogenizing in 20 mM HEPES, 0.25 M sucrose, 3 mM MgCl2, 0.2% IGEPAL CA-630, 3 mM β-mercaptoethanol, complete protease inhibitor tablet (Roche). Chromatin fragmentation was performed by sonication in 50 mM HEPES, 1% SDS, and 10 mM EDTA, using a Bioruptor (Diagenode) for 20 cycles of 30 s at the highest level. Cross-linked Chromatin was immunoprecipitated in 50 mM HEPES/NaOH at pH 7.5, 155 mM NaCl, 1.1% Triton X-100, 0.11% Na-deoxycholate, 1 mM EDTA, and complete protease inhibitor tablet, using 3 µg anti-ChREBP (Novus Bio NB400-135) antibody overnight. Antibodies were precipitated with a 33% slurry of precleaned protein A Sepharose beads in 0.5% BSA/PBS for 2 h. Crosslinking was reversed overnight at 65°C and DNA isolated using phenol/chloroform/isoamyl alcohol extraction. Enrichment of genomic sites in input and ChREBP ChIP was determined by qPCR, normalized to a site near the Ins gene, and evaluated using standard curves for each primer pair. The following primers were used: mIns -0.3 kb Fw: CTTCAGCCCAGTTGACCAAT; mIns -0.3 kb Rv: AGGGAGGAGGAAAGCAGAAC; mTxnip -0.2 kb Fw: CCGAACAACAACCATTTTCC, mTxnip -0.2 kb Rv: CGTGCACAGTTCTCCCATT; mPklr +0.23 Fw: CTCTGCAGACAGGCCAAAG; mPklr +0.23 Rv: TGCCAATGGAAGCCTTGTA; mKlf10 +1.3 kb Fw: GCATGTGAACAAAGCGTGAT; mKlf10 +1.3 kb Rv: TGCTCAGGAAGTAGGGGAAA. Two-Way ANOVA followed by Bonferroni post-test was used to analyze the data. padj < 0.05 was deemed significant.

### Correlation analyses

Hepatic cholesterol levels, gene expression, and clinical data were analyzed for correlations (Spearman) using the corrplot package. Correlations were deemed significant with a p-value < 0.05.

### Statical analysis

Samples were only excluded upon technical failure. Analyzes were performed in R studio (v. 4.2.1) or Prism software (v. 10.3.1). Specific statistical tests are described in each sub-section.

## Data availability

The transcriptome data from both in vivo and in vitro experiments are being deposited in the GEO database. Lipidomics data are being uploaded to the Metabolomics Workbench ^49^. Omics data will be available after publication.

## ARTIFICIAL INTELLIGENCE STATEMENT

The authors used Grammarly to improve readability and language, then reviewed and edited the content, taking full responsibility for the publication content.

## ACKNOWLEDGMENTS

This study was supported by grants of the German Research Foundation (DFG) to Oster, H. 353-10/1, GRK-1957, and CRC/TR 296 “LOCOTACT” (ID 424957847, TP13). The Metabolomics Workbench is supported by Metabolomics Workbench/National Metabolomics Data Repository (NMDR) (grant# U2C-DK119886), Common Fund Data Ecosystem (CFDE) (grant# 3OT2OD030544) and Metabolomics Consortium Coordinating Center (M3C) (grant# 1U2C-DK119889). de Assis, L.V.M. received a basic science research grant from the European Thyroid Association (ETA 2023) and support from the Knut and Alice Wallenberg Foundation as a Wallenberg Molecular Medicine Fellow. Metabolomics Workbench is supported by NIH grants U2C-DK119886 and OT2-OD030544.

## AUTHOR CONTRIBUTIONS

de Assis performed sample processing, data curation, and formal analysis. Inderhees processed and analyzed the lipidomics data. Wowro and Schupp handled ChREBP ChIP-PCR sample processing and data analysis. Heyde supported de Assis during diurnal sample collection and Fisher assisted with in vitro experiments. Jegodzinski, Affonso, Schönfels, Schenk, and Marquardt collected human samples and clinical data. Roßner conducted the histological analysis and scoring. The study was conceptualized by de Assis, Demir, and Oster. Oster provided supervision and secured funding. de Assis and Oster were responsible for the first manuscript draft. All authors reviewed and contributed to the manuscript revision.

## CONFLICT OF INTEREST

The authors declare no competing interests

---

**Figure S1:**
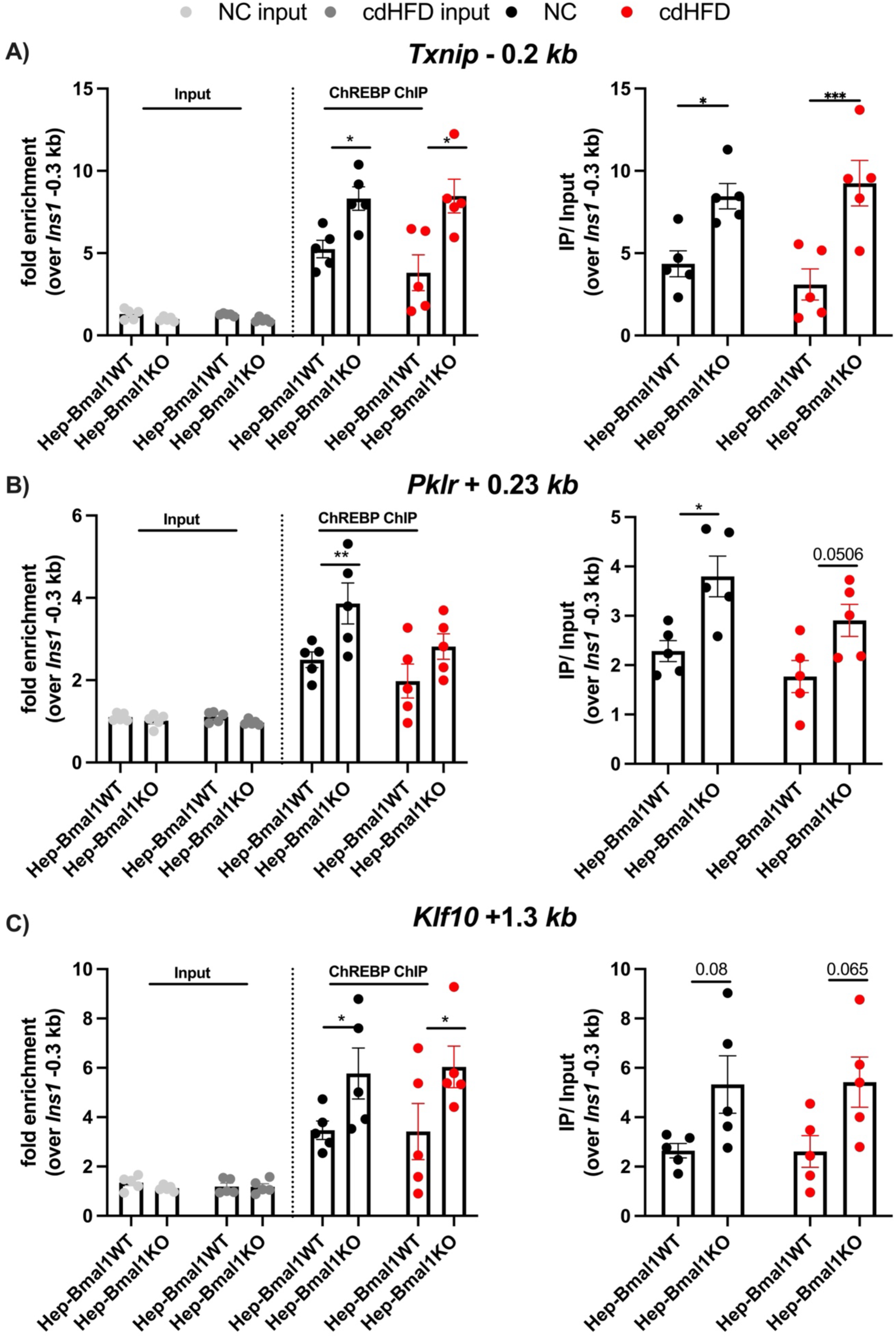
Chrebp ChipPCR in Hep-Bmal1WT and Hep-Bmal1KO livers. A – C) Analysis of selected *Chrebp* targets in livers of Hep-Bmal1WT or Hep-Bmal1KO mice fed with chow or cdHFD. N = 5 per group performed at ZT6.

**Figure S2:**
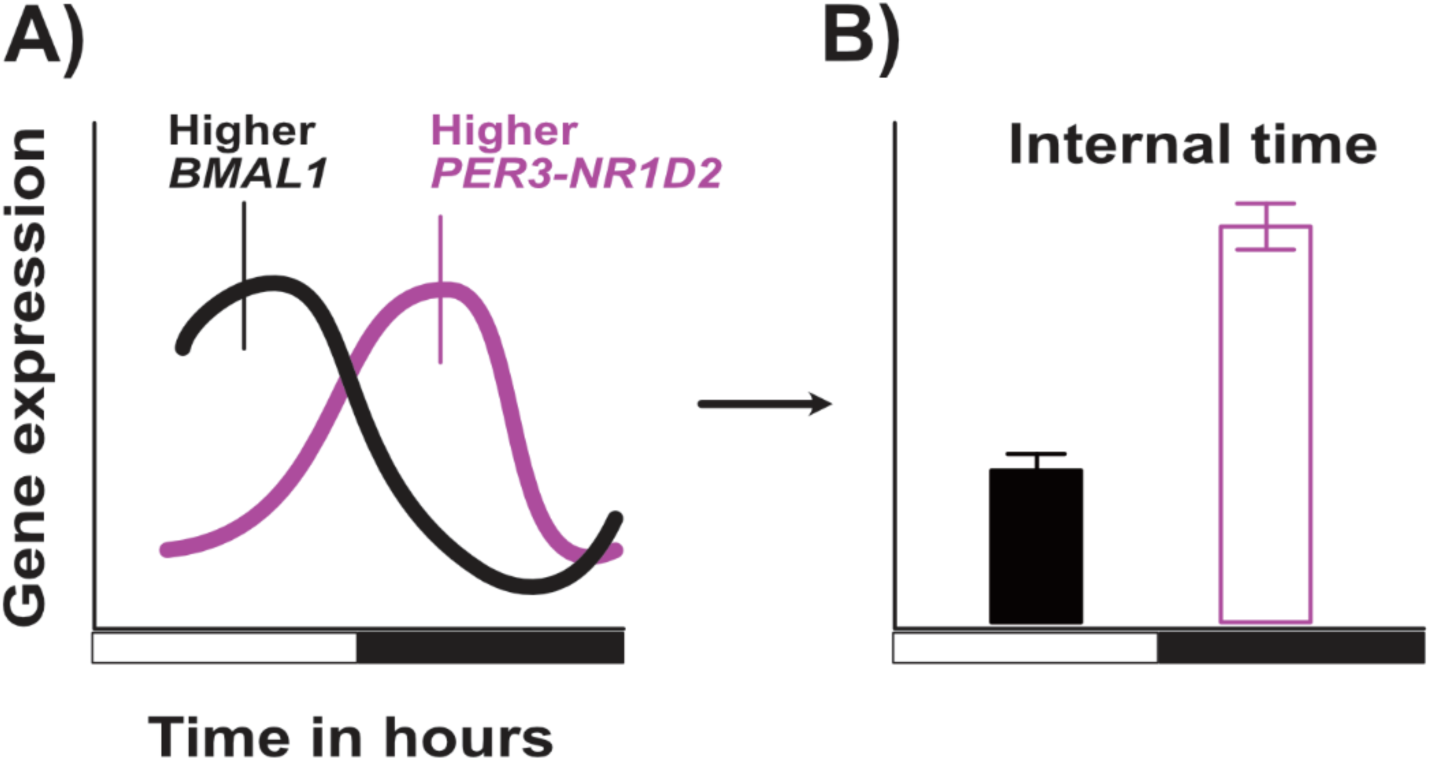
Defining the internal timing. A) Graphic representation of the diurnal gene expression of *BMAL1* (black curve) and *PER3*-*NR1D2* (purple curve). B) Presentation of the internal time calculation obtained from the graph described in A.

## Notes

### Competing Interest Statement

The authors have declared no competing interest.

